# Exosome derived multi-gene biomarker panel identifies the risk of liver metastasis in lung cancer patients

**DOI:** 10.1101/2023.12.12.571044

**Authors:** Kanisha A Shah, Rakesh M Rawal

## Abstract

**Background:** The lack of non-invasive methods for detection of early metastasis is a crucial reason for the poor prognosis of lung cancer (LC) liver metastasis (LM) patients. In this study, the goal was to identify circulating biomarkers based on a biomarker model for the early diagnosis and monitoring of patients with LCLM.

**Methods:** An 8-gene panel identified in our previous study was validated in CTC, cfRNA and exosomes isolated from primary lung cancer with & without metastasis. Further multivariate analysis including PCA & ROC was performed to determine the sensitivity and specificity of the biomarker panel. Model validation cohort (*n* = 79) was used to verify the stability of the constructed predictive model. Further, clinic-pathological factors, survival analysis and immune infiltration correlations were also performed.

**Results:** In comparison to our previous tissue data, exosomes demonstrated a good discriminative value with an AUC of 0.7247, specificity (72.48 %) and sensitivity (96.87%) for the 8-gene panel. Further individual gene patterns led us to a 5-gene panel that showed an AUC of 0.9488 (p = <0.001) and 0.9924 (p = <0.001) respectively for tissue and exosomes. Additionally, on validating the model in a larger cohort a risk score was obtained (RS >0.2) for prediction of liver metastasis with an accuracy of 95%. Survival analysis and immune filtration markers suggested that four exosomal markers were independently associated with poor overall survival.

**Conclusion:** We report a novel blood-based exosomal biomarker panel for early diagnosis, monitoring of therapeutic response, and prognostic evaluation of patients with LCLM.

## Introduction

Lung cancer is one of the most common tumors worldwide with high morbidity and mortality (1). Non-small-cell lung carcinoma (NSCLC), is the most prevalent subtype and patients with advanced NSCLC die within 18 months of diagnosis, with an overall five-year survival rate of only 15%. Lung cancer metastasis is the primary cause of death of the patients with liver being the most common metastatic organ for NSCLC. (2). Since there is a long incubation period from primary tumor to diagnosis of liver metastasis, many patients develop advanced liver metastasis at diagnosis (3). Moreover, these patients frequently develop metastasis to the other organs and lead to shorter survival and high mortality in patients with NSCLC liver metastasis. (4,5).

Histopathological evaluation using tissue biopsy from the site of the tumor along with radiological investigations are currently available diagnostic methods for lung cancer liver metastasis which is an invasive procedure. Some of the major limitations of invasive tissue biopsies are inadequacy of tissue, difficulty in accessing deep sites, and potential risk for the patient. (6). These currently available multistep, extended and invasive procedures are not unsuitable as a screening tool. Therefore, it is crucial to identify non-invasive, sensitive and specific circulatory biomarkers that will be extremely useful for screening and early detection of the disease (7). One of the promising approach to identify the potential biomarkers is to analyse the cancer-related biomolecules in bodily fluids. In addition, the non-invasive procedure for collection of blood makes biomarkers ideal as a screening tool for liver metastasis in lung cancer patients.

To identify novel metastasis-associated genes, our previous study detected differentially expressed profile of a multi gene panel specific for liver metastasis in primary lung tumor. The gene expression data acquired by real time PCR was further subjected to multivariate analysis like PCA and LDA which ultimately led us to the generation of a specific model (8). We found that out of the 5 models generated model 4 holds impressive potential for prediction of liver metastasis in primary lung cancer patients. Further, we also studied the individual sensitivity, specificity and cut-off values of each gene that were implicated in the model.

In this study, we validated 8 gene panel, in CTC, cfRNA and exosomes using real time PCR in primary lung cancer patients with & without metastasis and healthy controls, identified in our previous study in the tissue samples of the same cohort. ROC curve analysis was used to determine the sensitivity and specificity of the shortlisted gene panel and its correlation with the most accurate source of liquid biopsy component for early prediction of liver metastasis. Further DEP expression, clinic-pathological factors, PCA analysis and survival analysis correlations were performed. Using this model absolute quantification of candidate markers was done in exosomes from blood of primary lung cancer patients (n=79) resulting in the identification of potential biomarker panel with high sensitivity and specificity that could predict liver metastasis in high risk cohort.

## Materials & Methods

### Patient Characteristics

A total of 156 specimens from patients that passed the inclusion/exclusion criteria for the test and validation set were recruited in the study from December 2016 to July 2017. Peripheral blood (9 ml) was collected from patients with primary NSCLC with liver metastasis (n=32), primary NSCLC without liver metastasis (n=30) and healthy individuals (n=15) for the test cohort. For the validation cohort, peripheral blood (4 ml) was collected from primary NSCLC patients (n=79). Patients with only liver metastasis from primary lung cancer were included in the metastatic cohort and for the primary cohort only those patients with NSCLC subtype were included. Patients with multiple metastasis, HIV/HBsAg positive other malignant tumors or incomplete clinical data were excluded from the study. All the patients recruited in the test and the validation cohort in the study provided written consents for their sample procurement and use of them for research purpose.

### Culture of peripheral blood mononuclear cells for isolation of CTCs

Peripheral blood mononuclear cells (PBMNCs) were isolated by Ficoll-Hypaque density gradient centrifugation. The isolated PBMCs were cultured in RPMI growth media, and were incubated in a 5% CO2 incubator at 37 °C. After 24 h, the adherent cells were further expanded ex-vivo and observed daily for 30 days under a phase contrast microscope (Nikon Eclipse TS100). Adherent cells that appeared CTC-like were counted manually using a microscope to calculate the number of cells present in each cultured sample.

### Tumor Sphere formation & Cytotoxicity assay

Cells were plated to 6 well ultra-low attachment plates (Corning; New York, USA) at a density of 5x 10^3^ cells/well and further expanded with RPMI-1640 supplemented growth media. Wells containing spheres were counted manually under inverted phase contrast microscope and the percentage of cells exhibiting sphere-forming capacity was calculated by dividing the number of spheres by the number of cells seeded per well. For the cytotoxicity assay the isolated CTCs were seeded at a density of 1x10^4^ cells/well in 96-well plate and exposed to Cisplatin and Carboplatin drugs for 24 hours in increased concentrations ranging from 0.25 to 1.0 µg/ml. Further, 5 mg/ml MTT (Hi-Media, India) and 100 µl of DMSO were added to each well for formazon crystal formation and the cells were incubated at 37 °C. Absorbance was taken at 590 nm with a reference filter of 620 nm using an ELISA reader (Multiskan Spectrum Microplate Reader, Thermo Scientific). All data were normalized against DMSO blank controls.

### Flow cytometry

The purity of cultured CTC subpopulation was validated by gating the sorted cells with conjugated primary antibodies of CD44-FITC (Stem Cell Technologies), CD24-PE, CK-FITC and CD45-PE (BD Biosystems, USA) using flow cytometry analysis and for characterization of exosomes, exosome-coated beads (Stem Cell Technologies; USA), were stained with primary mouse monoclonal antibodies directed against CD63, CD81 and CD9. The percentage of each population was calculated by acquiring the sample in FACS Canto II instrument and result was analyzed using FACS Diva software (BD Biosystems, USA).

### Exosome Isolation & Characterization

Exosomes were isolated from 1 ml of serum using a Total Exosome Isolation Kit (Invitogen, Thermo Fisher Scientific Inc., USA), according to the manufacturers protocol. Exosomal pellets were split and resuspended (i) in 300 µl PBS for characterization, (ii) in 500 µl PBS for RNA isolation and real time PCR and the remaining exosomes were stored at −80 °C. The purified exosomes were fixed with 2% (w/v) para-formaldehyde (PFA) in PBS (pH 7.4); dropped onto a carbon-coated grid and left to dry at room temperature. Further they were fixed with 1% (w/v) glutaraldehyde and stained with saturated aqueous uranyl oxalate. Exosomes were then re-suspended in a solution of 0.4% (w/v) uranyl acetate & 1.8% (w/v) methylcellulose, incubated on ice, excess liquid was drained off and the grid was air dried. Exosomes were viewed using a Field Emission Scanning Electron microscope (JSM-7600F; JEOL, Tokyo, Japan) (9). Nanoparticle Tracking Analysis (NTA) was performed using a Nanosight LM10 and NTA 2.3 Software (NanoSight, Wiltshire, UK). EV samples were resuspended in filtered PBS. Three 60-sec videos were recorded of each sample with camera level and detection threshold set at 10. Temperature was monitored throughout the measurements.

### Quantitative Real Time PCR (qRT-PCR)

Total RNA from CTCs was isolated using RNeasy kit (Qiagen 74106) following manufacturer’s instructions whereas TRIzol reagent (Invitrogen, USA) was used for total RNA isolation from exosomes and cell free RNA from serum of patients and healthy individuals. 1 µg total RNA was reverse transcribed using the cDNA archive kit (Applied Biosystems – ABI, USA) following manufacturer’s instructions and the resulting cDNA was used as a template for Real Time PCR analysis. qRT-PCR was performed using AriaMX (Agilent Technologies, USA) with a SYBR Green PCR Master Mix (Primer sequences are provided in Supp Table I). The generation of a specific PCR product was tested using the ΔΔCT method with β-actin as the endogenous control and average C_T_ value was calculated for the quantification fold change analysis.

**Table I:** Correlation between tissue gene panel expression levels and clinical characteristics.

| Sr. No |  | n values (%) | CXCL12 (Mean rank values) | p | TGFBR2 | p | CD44v6 | p | HIF1A | p | KRT7 | p |
| --- | --- | --- | --- | --- | --- | --- | --- | --- | --- | --- | --- | --- |
| 1 | Age | 33 |  | 0.417 |  | 0.857 |  | 0.259 |  | 0.787 |  | 0.1074 |
|  | >=59 | 18 | 15.72 |  | 16.69 |  | 15.25 |  | 16.56 |  | 14.5 |  |
|  | <59 | 15 | 18.53 |  | 17.37 |  | 19.1 |  | 17.53 |  | 20 |  |
| 2 | Habit |  |  | 0.096 |  | 0.689 |  | 0.253 |  | 0.0818 |  | 0.624 |
|  | Smoking | 23 | 18.73 |  | 17.44 |  | 18.3 |  | 15.04 |  | 16.43 |  |
|  | Non-smoking | 10 | 12.39 |  | 15.83 |  | 14 |  | 21.5 |  | 18.3 |  |
| 3 | Differentiation |  |  | 0.048 |  | 0.685 |  | 0.671 |  | 0.896 |  | 0.659 |
|  | poorly diff | 12 | 12.58 |  | 16.58 |  | 16 |  | 16.67 |  | 15.91 |  |
|  | diff unknown | 21 | 19.52 |  | 17.24 |  | 17.57 |  | 17.19 |  | 17.55 |  |
| 4 | Stage |  |  | 0.681 |  | 0.96 |  | 0.008 |  | 0.857 |  | 0.303 |
|  | III | 15 | 17.88 |  | 17.14 |  | 21.8 |  | 16.58 |  | 14.81 |  |
|  | IV | 18 | 16.42 |  | 16.89 |  | 13 |  | 17.27 |  | 18.42 |  |
| 5 | EGFR Status |  |  | 0.275 |  | 0.322 |  | 0.427 |  | 0.645 |  | 1 |
|  | wildtype | 12 | 19.64 |  | 14.59 |  | 15.21 |  | 15.86 |  | 17.05 |  |
|  | mutation | 21 | 15.68 |  | 18.2 |  | 18.02 |  | 17.57 |  | 16.98 |  |
| 6 | AE1 |  |  | 0.417 |  | 0.303 |  | 0.899 |  | 0.528 |  | 0.496 |
|  | Positive | 13 | 15.27 |  | 19.19 |  | 17.31 |  | 18.35 |  | 18.46 |  |
|  | Negative | 20 | 18.12 |  | 15.58 |  | 16.8 |  | 16.12 |  | 16.05 |  |
| 7 | Chromogranin |  |  | 0.818 |  | 0.896 |  | 0.043 |  | 0.802 |  | 0.603 |
|  | Positive | 11 | 16.56 |  | 18.28 |  | 13.45 |  | 17.47 |  | 17.94 |  |
|  | Negative | 22 | 17.41 |  | 17.76 |  | 20.77 |  | 16.56 |  | 16.12 |  |
| 8 | CEA |  |  | 0.031 |  | 0.96 |  | 0.735 |  | 0.0093 |  | 0.0423 |
|  | Positive | 15 | 21 |  | 16.87 |  | 17.63 |  | 21.83 |  | 12.64 |  |
|  | Negative | 18 | 13.67 |  | 17.11 |  | 16.47 |  | 12.97 |  | 19.5 |  |
| 9 | CK20 |  |  | 0.541 |  | 0.638 |  | 0.696 |  | 0.1556 |  | 0.483 |
|  | Positive | 6 | 14.75 |  | 18.75 |  | 15.75 |  | 22.17 |  | 19.58 |  |
|  | Negative | 27 | 17.5 |  | 16.61 |  | 17.4 |  | 15.85 |  | 16.43 |  |
| 10 | CK7 |  |  | 0.173 |  | 1 |  | 0.643 |  | 0.757 |  | 0.152 |
|  | Positive | 28 | 17.98 |  | 17.02 |  | 16.64 |  | 17.09 |  | 18.04 |  |
|  | Negative | 5 | 11.5 |  | 16.9 |  | 19 |  | 19.9 |  | 11.2 |  |
| 11 | CDX2 |  |  | 0.218 |  | 0.794 |  | 0.899 |  | 0.332 |  | 0.238 |
|  | Positive | 12 | 14.62 |  | 16.38 |  | 17.31 |  | 19.21 |  | 19.67 |  |
|  | Negative | 21 | 19.02 |  | 17.36 |  | 16.8 |  | 15.74 |  | 15.48 |  |
| 12 | TTF-1 |  |  | 0.033 |  | 0.802 |  | 0.708 |  | 0.322 |  | 0.158 |
|  | Positive | 15 | 20.97 |  | 17.5 |  | 16.3 |  | 15.13 |  | 14.37 |  |
|  | Negative | 18 | 13.69 |  | 16.58 |  | 17.58 |  | 18.56 |  | 19.19 |  |

### Principal component analysis

To obtain a reliable model for prediction of liver metastasis, data was analyzed statistically by applying Principal Component Analysis (PCA) method. Data dimensionality was reduced by using an orthogonal transformation to convert correlated variables into uncorrelated variables, which are termed principal components. PCA score was obtained using the following formulae:

PCA score = C1V1 + C2V2 + C3V3 + C4V4 + + CnVn

where C1, C2, C3 … Cn are coefficients of each of the variables and V1, V2, V3 … Vn are the values of original variables

The accuracy, specificity and sensitivity of the PCA based model was securitized to check for the robustness of the model. For this, patient stratification process was simulated 100 times by resampling. The patients were divided into primary lung cancer set (30 patients, 48%) and liver metastatic test set (32 patients, 52%) groups. PCA was applied to the data sets to determine the coefficient of each variable, and a PCA score was generated for each patient. The data sets were then stratified into groups based on the mean value of the PCA scores. Statistical analyses and the PCA algorithm were performed using SPSS 20.00 software and R and a p-value of < 0.05 was considered statistically significant.

### Linear discriminant analysis & receiver operating characteristics curve analysis

Linear Discriminant Analysis (LDA) was used to determine whether mRNA expression patterns could accurately discriminate liver metastasis in an independent data set. The accuracy of the predicted model was calculated using 1000 repetitions of a random partitioning process to regulate the number and proportion of false discoveries (10). For diagnostic accuracy and discriminating metastatic tumor from primary tumor, a held-out test set from each database was utilized to evaluate the performance of each of the different classifiers. Receiver operator characteristics curves (ROC) were generated and AUCs of each classifier were calculated using MedCalc (Belgium, Europe). To understand the false positives and/or weaknesses of our classifiers, images frequently misclassified by the classifiers were also reviewed.

### Connections between liver metastasis-associated genes, immune cell infiltration and immune checkpoints

TIMER (https://cistrome.shinyapps.io/timer/) is a computational tool that aids in comprehensively exploring the molecular characterization of tumor immune interactions across diverse tumors (11). It calculates the abundances of six immune infiltrates (CD8 + T cells, CD4 + T cells, B cells, neutrophils, macrophages, and dendritic cells) based on RNA-Seq expression profiles data. We used TIMER to evaluate correlations between liver metastasis-associated gene expression and six immune cells and CTLA4, CD274, PDCD1, & PDCD1LG2 checkpoint genes.

### Statistical Analysis

All experiments were reproducible and each set was repeated at least three times to check the reproducibility. All data were recorded as mean ± standard deviation (SD) unless stated otherwise. PCA and LDA were performed using SPSS 20.00 statistical software (Chicago, IL, USA). Kaplan Meier Survival curve analysis was performed using Graph Pad Prism 7.0 software (GraphPad Software Inc.). Chi Square test and Mann Whitney U test were performed using the SPSS 29 software (IBM, USA) for clinical correlation with differential gene expression data. P-value < 0.05 was considered to be statistically significant

## Results

### Isolation and characterization of CTCs from PBMNCs of primary and metastatic tumors

PBMCs were isolated from peripheral blood of NSCLC patients with and without liver metastasis and cells were expanded ex-vivo after incubation for 24 h. The cells were cultured for a period of 15-20 days and circulating tumor stem like cells were observed as circular cells with spikes on their circumference. These cells were considered to be CTCs and their morphology and other characteristics were examined by phase contrast microscopy. Furthermore, we characterized these CTCs to check the purity by analyzing the presence of CSC and epithelial markers like CD44, CD24, CD45 and CK using flow cytometry. Based on the flow cytometry analysis, we identified three distinctive sub-populations: (i) CD44+ CD24-; (ii) CD44+ CK7+; (iii) CK7+ [Supp Fig I(A)] for primary lung cancer patients whereas the other hand, CTCs isolated from patients with lung cancer liver metastasis exhibited three distinctive subpopulations: (i) CD44+ CD24+; (ii) CD44+ CD24-; and (iii) CD24+ [Supp Fig I(B)]. However, absence of CD45+ cells in the sorted population confirmed the non-lymphocytic characteristic of these cancer stem like cell population.

### Increased self-renewal potential and drug resistance properties of CTCs

Intrinsic sphere forming ability of the isolated CTCs was assessed and they demonstrated enhanced colony forming ability with a significant increase in the number and size of spheres of metastatic patients [Supp Fig I(D)] as compared to CTCs isolated from primary tumors [Supp Fig I(C)]. Further, on assessing the proliferation rate, CTCs isolated from both the primary and metastatic patients demonstrated a colony forming efficiency of nearly 60-70% with the colonies exponentially increasing at regular intervals, suggestive of the fact that both the subpopulations possess the intrinsic self-renewal potential but the metastatic subpopulation had a 15% higher proliferation efficiency as compared to the other counterpart [Supp Fig I(E) & (F)].

Drug resistance is an intrinsic property of CSC population as these slow growing cells tend to often escape the conventional therapeutic regimen unlike the proliferating cells. Thus, we assessed the cytotoxic effect of carboplatin and cisplatin on CTCs isolated from primary lung cancer with and without metastasis. The CTCs from lung cancer and liver metastasis demonstrated 88.2% and 89.32 % viability respectively as compared to merely 36.43% viability of MNC cells from healthy individuals even at higher concentrations of Cisplatin (100 µM; p-value < 0.001) Moreover, lung cancer derived CTCs showed viability of 85% and those of metastasis demonstrated a viability of 88% at higher concentrations of Carboplatin (100 µM; p-value < 0.001) as compared to the healthy MNC cells (viability 41.8%) [Supp Fig I(G) & (H), Supp Table I & II]. Collectively, these results are suggestive of the fact that the CTCs were significantly chemo-resistant and showed dose dependent drug resistance to advanced chemotherapeutic drugs - Cisplatin and Carboplatin.

### Characterization of the exosomes derived from primary lung cancer cells with and without liver metastasis

To investigate the role of exosomes in lung cancer liver metastasis, exosomes were isolated from the serum of lung patients with and without liver metastasis and their identity was confirmed by electron microscopy, flow cytometry and NTA. The vesicles isolated from the serum appeared as round-shaped 40 - 50 nm diameter vesicles under electron microscopy [Supp Fig II(A)]. We further confirmed whether these vesicles were exosomes by performing NTA analysis [Supp Fig II(B)]. Results of flow cytometry also showed CD63 and CD81to be present in exosomes derived from serum of lung cancer patients with and without liver metastasis which is are commonly used exosomal markers [Supp Fig II(C)].

### Differential gene expression analysis in CTC, Exosomes and cfRNA: Identifying a better marker for liquid biopsy

Quantitative gene expression patterns of CXCL12, CK7, CDH1, CTNNB1, HIF1A, MUC16, TGFBR2 and CD44v6 were analyzed from CTC, exosomes and cell free RNA of lung cancer patients with and without liver metastasis. CXCL12, CK7, CDH1, CTNNB1, TGFBR2 and CD44v6 were upregulated in CTC as well as cfRNA [Fig I(A) &(C)] whereas all these genes were downregulated in exosomes of primary lung cancer with and without metastasis [Fig I(B)]. Interestingly, it was observed that HIF1A was downregulated in primary lung cancer but was upregulated in liver metastasis in CTCs and exosomes [Figure I(A) & (B)]. Moreover, MUC16 was only expressed in CTCs derived from liver metastasis but were not expressed in CTCs of primary lung cancer [Fig I(A)]. In exosomes, it was seen that MUC16 was upregulated in primary lung cancer whereas it was downregulated in liver metastasis [Figure I(B)]. In cfRNA the expression of all the genes were observed to be upregulated in both the cohorts [Figure I(C)].

### Generation of Candidate Biomarker Panel

ROC analysis, was performed, using the values, to determine and confirm robustness of the models generated using different sample source (Exosomes, cfRNA and CTC) comprising of the genes (CD44v6, CXCL12, CTNNB1, CK7, HIF1A, TGFBR2, MUC16, CDH1) that were shortlisted for the tissue model [Fig I(D)] (8). Area under the curve (AUC) specificity and sensitivity are mentioned in Supp Table II. In order to determine if the multigene signature has any practical application, we performed ROC analysis for individual genes as seen in Supp Fig III and the genes with maximum significant sensitivity, specificity & AUC in exosomes, cfRNA and CTC were further considered for the final biomarker model. From the ROC curve analysis of individual genes, we further speculated that a more precise model for prediction of lung liver metastasis could be formed with 5 genes (CXCL12, CK7, CD44v6, HIF1A, and TGFBR2) algorithm as described in [Fig I(E)] and Supp Table III. Moreover, it was observed that exosomal model came out as the best model and was at par with the tissue model and hence can be used for predicting liver metastasis non-invasively with the highest AUC of 0.975 a Specificity and Sensitivity of 90% and 96.87% respectively showing good discriminative ability between primary and metastatic tumors. CTC and cfRNA did not show significant AUC, sensitivity and specificity as a diagnostic model and hence cannot be used for early prediction of liver metastasis in lung cancer patients.

### PCA Analysis distinguishing between primary and metastatic carcinomas

The liver is one of the most frequent sites of metastasis. Distinguishing between primary and metastatic carcinomas is important due to differences in their management. Therefore, genes from FNAC samples were profiled and added to the primary lung carcinoma dataset. The metastases originated from lung carcinomas. When the combined dataset was analyzed by PCA, primary and metastatic adenocarcinomas were clearly distinct in the tissue and the exosome cohorts [Fig II(C) & (D)]. The small sample size for metastatic adenocarcinomas precluded us from performing class prediction analysis after training-to-test set allocation. Instead, we performed cross-validation analysis using all samples as a training set. The permutation *p*-values for the misclassification rate (at feature selection *p*<10^−6^) were CCP: 0.02, LDA: 0.01, 1-NN: <0.01, 3-NN: 0.01, NC: 0.01 and SVM: <0.01. In primary *vs.* metastatic carcinomas, *CXCL12, CD44v6, CK7, HIF1A and TGFBR2* were differentially expressed at *p*<10^−6^; the level of expression of all the above-mentioned genes from exosomes of metastatic carcinoma were downregulated as compared to tissue samples where CD44v6 was upregulated and the other genes had a downregulated expression.

### Risk Score equation for lung cancer liver metastasis

(0.713*0.47) +(0.570*0.340) +(0.632*0.346) +(0.774*10.305) +(0.588*0.468) = 8.689686

### Risk Score equation for primary lung tumor

(-0.711*0.076) +(0.927*0.043) +(0.908*0.035) +(0.302*0.019) +(0.791*0.166) = 0.154649

When we further performed the ROC for the model we had got a cut off value of >0.2 for predicting metastasis (mentioned in Supp Table III). Our results comply with the associated criteria and thus from the above equation we can say that these markers together can be used for predicting liver metastasis in lung cancer patients. Moreover, from the above equation we observed that the metastasis patients have an average value of 8.689686 which is higher than the cut off value whereas for the primary patients the value obtained was 0.154649 which is lower than the cut off value for prediction. The combination variables had the best overall accuracy, resulting in an AUC of 0.80 (95% CI = 0.74 to 0.86). Next, ideal cutoff points were assessed as maximum sum of sensitivity and specificity. This revealed both a sensitivity and a specificity of 98.2% and 95.29% respectively and the AUC of 0.992 (95% CI = 0.968 to 0.996).

### Gene Expression and Clinical/Pathological Correlation in metastatic tissue and exosome cohorts

We further analysed the correlation between CXCL12, CD44v6, HIF1A, TGFBR2 and KRT7 expression in metastatic tumor tissues & exosomes with sex, age, habit, Tumor Differentiation grade, TNM stage, EGFR mutation status, AE1, Chromogranin, CDX2, CEA, CK20, TTF-1 & CK7. Results showed the correlation between CXCL12 and differentiation grade, CEA and TTF-1 protein expression, CD44v6 was associated with tumor stage, chromogranin expression whereas HIF1A and KRT7 were significantly associated with CEA protein expression in the metastatic tissues. On the other hand, in the exosome cohort it was observed that CD44v6 was associated with stage, Chromogranin, CK20 & TTF-1 protein expression, CXCL12 was significantly correlated to EGFR mutations, TGFBR2 was significantly associated with AE1 protein expression and HIF1A with CEA protein expression (p value < 0.05) (Table I & II).

**Table II:**
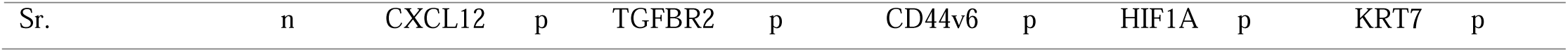
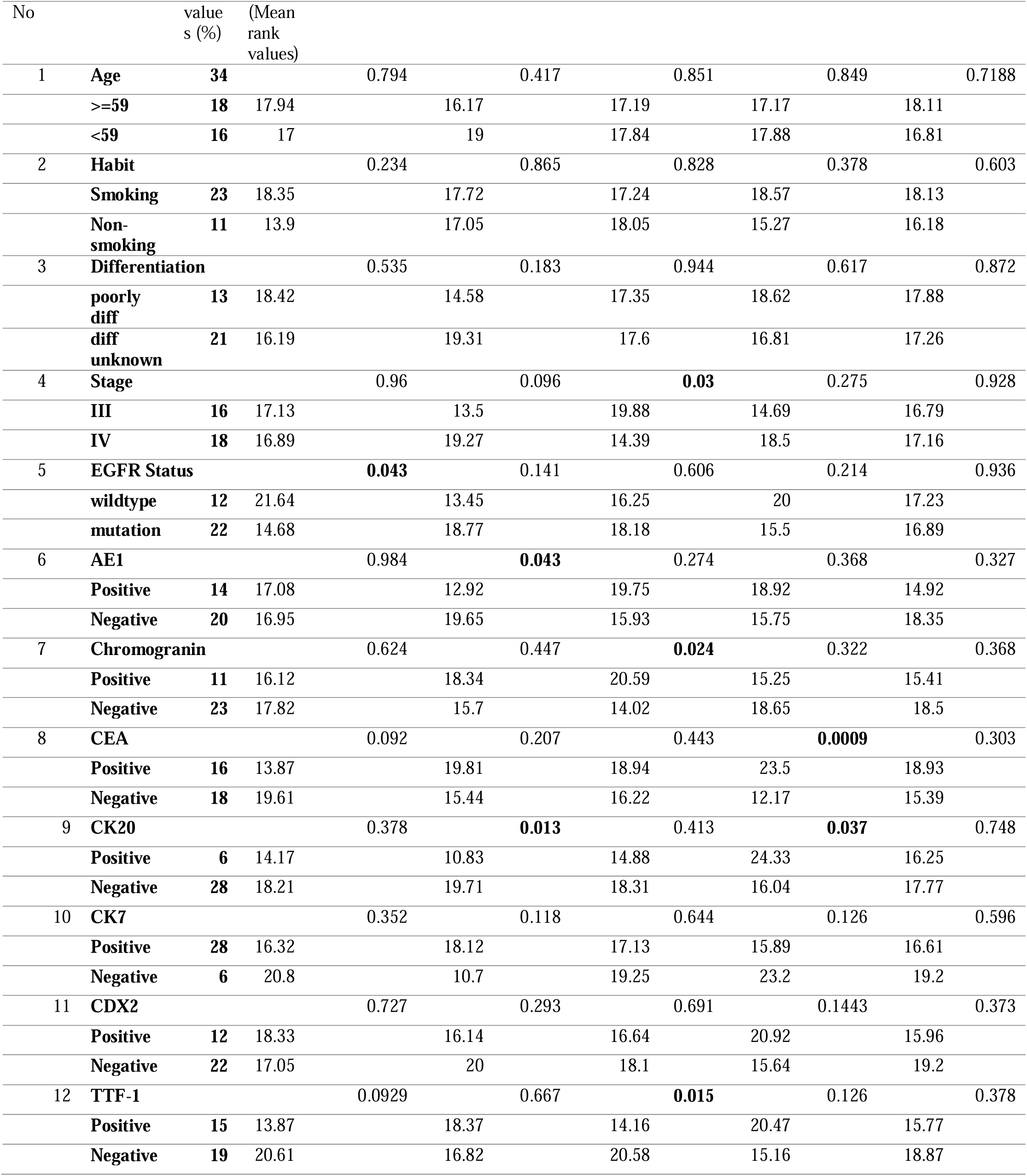
Correlation between exosome gene panel expression levels and clinical characteristics.

### Clinical Validation, Risk Stratification and Heatmap generation

The five-marker signature was tested in the exosomes isolated from the blood of patients in the validation set (n= 79). The score plot demonstrated that patients at high-risk had poorer outcomes and developed liver metastasis as compared to patients at low-risk. Figure shows the distribution of the Risk Score, and heatmap of the 5-gene signature in the validation set [Fig III (A) & (B)]. It was observed that out of 79 patients 44 patients had a high risk for developing liver metastasis as compared to the other 35 patients. In the high-risk cohort, it was observed that out of 44 patients 39 developed liver metastases, 3 patients developed multiple metastasis (brain, bone & liver), whereas 2 patients did not develop any distant metastasis. In the other 35 patients 9 patients could not be evaluated at follow up as there was no data available whereas the other 26 did not develop any distant metastasis. The survival prognosis was significantly worse for the high-risk group as compared to the low risk cohort [Fig III]. These results demonstrate that the algorithm developed could accurately predict liver metastasis in 42 out of 44 patients (95% accuracy) with high-risk score based on the equation derived.

### Clinical relevance and Kaplan–Meier survival analyses of the metastasis associated genes in tissue and exosome cohorts

To further demonstrate the clinical significance of the 5-gene signature in patients with lung cancer liver metastasis, the association between gene expressions and various clinico-pathological variables was investigated by real-time quantitative PCR in 32 lung cancer liver metastasis patients. We found that in the tissue cohort CXCL12 expression was strongly correlated with differentiation, CEA & TTF-1 protein expression, CD44v6 was associated with tumor stage and chromogranin protein expression and HIF1A & KRT7 showed positive correlation with CEA expression alone (Table I). In the exosome cohort CXCL12 expression was strongly correlated with EGFR status, CD44v6 was associated with tumor stage, TTF-1 expression & chromogranin protein expression and HIF1A & TGFBR2 showed positive correlation with CEA expression & CK20 expression (Table II). The Kaplan–Meier survival analyses of the metastasis associated genes in lung cancer patients with liver metastasis was evaluated. Among all the patients, low expression of KRT7 (*P* = 0.0079) & TGFBR2 (*P* = 0.0075) exhibited significantly worse Overall Survival (OS) compared to high expression in tissue cohort [Fig II (A)]. Contradictorily it was observed that low expression of KRT7 (*P* = 0.0009), TGFBR2 (*P* = 0.024), & HIF1A (*P* = 0.0246) and high expression of CD44v6 (*P* = 0.0326) was associated with worse OS in the exosome cohort [Fig II(B)].

### Association of marker panel genes to immune cell infiltration in LCLM

To ascertain if correlations existed between tumor infiltrating immune cells and metastasis-associated gene levels, we examined associations using TIMER. KRT7 expression was positively correlated with infiltrating CD8 + T cells (*P* = 0.00923), B cells (*P* = 6.50e-10), and myeloid dendritic cells (*P* = 0.0048). Similarly, HIF1A was also positively correlated with infiltrating B cell (*P* = 0.03), CD8 + T cells (*P* = 8.14e-08), macrophages (*P* = 7.32e-07), neutrophil (*P* = 3.66e-18), and myeloid dendritic cells (*P* = 4.24e-09). Moreover, CD44, CXCL12 and TGFBR2 were positively correlated with all tumor infiltrating immune cells [Fig IV]. As current immunotherapy strategies rely on immunological checkpoint inhibitors (12, 13) we used TIMER to investigate co-expression relationship of all 5 genes with immune checkpoint-related genes. CD44, CXCL12 and HIF1A displayed strong co-expression relationships with CD274, CTLA4, PDCD1, and PDCD1LG2, whereas KRT7 had co-expression relationships with CTLA4, & PDCD1LG2 and TGFBR2 has co-expression with CTLA4, CD274 and PDCD1LG2 [Fig V].

## Discussion

Non-small cell carcinoma (NSCLC) is one of the major causes of cancer death globally. Liver metastasis is one of the most critical prognostic factor for NSCLC (14). Early detection of heterochronous lung cancer liver metastasis, may provide useful information for designing treatment strategy to improve patient survival. Most patients with early lung cancer have a 70% survival rate after appropriate surgical resections, yet many patients already develop distant metastasis, which is difficult to detected solely by imaging methods (15). Moreover, patients with poor tumor differentiation and higher TNM stage develop distant metastasis, and their average survival period is 4 months (16). However, TNM staging is unable to facilitate accurate prognosis and hence it becomes difficult to determine treatment. It has been recommended that the traditional TNM staging should include number of involved metastatic sites, number of metastatic foci per involved site and the diameter of each metastatic focus. This will help in prediction of metastasis for more accurate tumor staging, which could improve treatment efficacy and increase survival rates (17).

Presently, serum biomarkers including sCEA, sCA125, sCA199 and sNSE have been widely used in the diagnosis and monitoring the progression of lung cancer liver metastasis (18). However, the sensitivity of CEA and CA199 have always been lower compared to that of sCA125 and NSE. Unfortunately, the results of predictive ability of these markers or in combination with other indices are greatly discrepant (19, 20). Therefore, identification of non-invasive biomarkers for early diagnosis of NSCLC LM and dynamic monitoring of disease status is a key research imperative. There is a growing consensus that biomarker panels have higher specificity and sensitivity than single biomarkers and may be more effective in detecting cancer (20, 21). Herein, we sought to discover the unique patterns of circulating markers in NSCLC patients with LM, and to identify biomarkers with sufficient sensitivity and specificity for use as a supplement or substitute for invasive biopsy in clinical practice. In the present study, exosomal CXCL12, TFBR2, CD44v6, HIF1A and KRT7 were identified as novel diagnostic biomarkers for NSCLC LM. Moreover, the combination of this marker panel may be a sensitive biomarker for follow-up monitoring of therapeutic response, as well as a prognostic marker in patients with LM.

In this study, we used a previously established multi-gene panel for generating a model specific for prediction of liver metastasis. We generated the model using PCA and LDA and further calculated the accuracy of the model generated by ROC curve analysis that showed accuracy greater than 90%. Additionally, the Kaplan Meier survival data also showed a significance (p<0.0001) amongst the gene expression patterns of the different genes and their association with overall survival (OS) in metastatic patients. Moreover, the developed equation with the co-efficient and the risk score aided in predicting metastasis in the primary lung cancer validation cohort with an accuracy of 92%. Furthermore, there was a significant association between the gene panel expression and the tumor infiltrating markers signifying the important role of these biomarkers in the development and diagnosis of liver metastasis. Thus, the five-gene panel (CXCL12, KRT7, CD44v6, TGFBR2 and HIF1A) can be a highly significant predictor of liver metastasis outcome independent of the standard prognostic criteria. Additionally, the ability of this panel to accurately predict recurrence in the liver independent of stage of the primary tumor is likely to be a useful enhancement to routine staging.

The biomarker panel identified in this study are not conventional tumor-derived cancer biomarkers but their combination generating a cut off score that aids in early detection of liver metastasis. This panel shows better diagnostic accuracy in comparison to the individual markers. Stromal cell-derived factor 1 (SDF-1) also known as CXCL12 is a crucial chemotactic factor that stimulates proliferation, adhesion, dissociation, migration, survival of tumor cells and the formation of tumor-associated vessels and invasion in a wide variety of tumor cells, especially reported in liver metastasis (22). Furthermore, the CXCL12/CXCR4 has been recognized as a prognostic marker in different cancers and preclinical models; signifying that metastasis is mediated by CXCR4 activation and migration of tumor cells towards CXCL12 expressing organs. Moreover, KRT7 is a membrane-cytoskeletal linker which contributes to regulation of cell adhesion and overexpression is associated with cellular transformation. KRT7 plays an important role in preserving tissue architecture, tumor progression, invasion, and metastasis. Cytoskeletal component levels, such as the degree of keratinization, have been reported to be valuable in identifying patients at risk for developing regional metastases (23). CD44 transmembrane glycoproteins are cell adhesion molecules that have been associated with aggressiveness and metastasis. Through HIF-1/2α, hypoxia transcriptionally controls a host of factors that contribute to increased resistance to radiation and chemotherapy, and to the emergence of a more aggressive phenotype by regulating CD44v6 in metastasis (24–27) The functional effects of the upregulation of CD44v6 and its variant isoforms under hypoxic conditions should be considered, since cell signaling events that promote anchorage-independent tumor cell growth, survival, migration and metastasis occur through the binding of hyaluronan with CD44 (28). Upregulation of both CD44 and hyaluronan under hypoxic conditions most likely amplify the signaling events and lead to tumor progression (29). Hypoxia Inducing Factor 1α (HIF1A) plays an important role in the formation of liver metastasis. Further studies have reported that HIF1A overexpression enhances ZEB1 trans-activity by binding to its promoter leading to a loss in E-cadherin causing increased invasion, migration and tumor progression (30). Lastly, TGFB signaling is also suggested to be involved in the malignant progression of tumors and can induce epithelial–mesenchymal transition and promote tumor cell invasion, and may also have angiogenic and immunosuppressive effects on the tumor microenvironment, all of which promote metastasis.<u>8</u> It has been observed that reduced TGFBR2 expression significantly is correlated with the aggressive features in Hepatocellular carcinomas as well as other tumors. Morover, TGFBR2 has also been reported in liver metastasis arising from colorectal tumors (31). Additionally, all these markers have an important role in liver metastasis and hence the generated model will assist in the early diagnosis of liver metastasis.

Although combination of exosomal serum levels of CD44v6, HIF1A, TGFBR2, KRT7 and CXCL12 seem to be promising biomarkers and may guide in the development of novel targeted drugs, some limitations of our study should be considered while interpreting the results. For example, there might be some inherent biases since clinical parameters are variable between institutions and individual clinicians. Therefore, a well-designed and large-scale multicenter follow-up cohort study is warranted to provide more robust evidence.

## Conclusion

In summary, using a multiplexed, molecularly driven approach, we have identified a panel comprising CXCL12, KRT7, CD44v6, TGFBR2 and HIF1A that can predict recurrence in the liver independent of conventional prognostic criteria and identify patients with lung cancer who will develop liver metastasis despite undergoing definitive surgery and/or treatment. Increasing numbers of alterations in these genes predict poorer prognosis. As an outcome of this study, we report a sensitive biomarker panel which can be developed into a multi-parameter rapid testing kit to explore its potential in clinical settings. However, the study being a pilot study in nature, the outcome needs to be evaluated in a larger patient population. For future directions, cross-validation of this prediction model in terms of accuracy, precision, sensitivity, specificity and positive and negative predictive values using a separate large cohort of lung cancer liver metastasis cases, disease controls and healthy controls is needed so that the potential value of this prediction model as a biomarker panel in the clinical setting can be explored and a rapid detection kit can be developed for the model to facilitate population screening.

## Supporting information

Supplementary Figure III

## Data Availability

All data generated or analysed during this study are included in this article and its supplementary material files. Further inquiries can be directed to the corresponding author

## Contributions

KS collected the data. KS and RMR designed the research study. KS and RMR analysed the data and drafted the manuscript. RMR proofread the final manuscript. All authors approved the final version of the manuscript.

## Acknowledgement

I would like to achnowlege DHR – ICMR for my current project as a Young Scientist (File No: File No.R.12014/52/2022-HR)

## Informed consent

Written Informed consent was obtained from all individual participants included in the study

## Ethical approval

All procedures performed in studies involving human participants were in accordance with the ethical standards of the institutional and/or national research committee [The Gujarat Cancer & Research Institute (EC-O-92-2014) and Gujarat University (GU/IEC/07/2018)] and with the 1964 Helsinki declaration and its later amendments or comparable ethical standards.

---

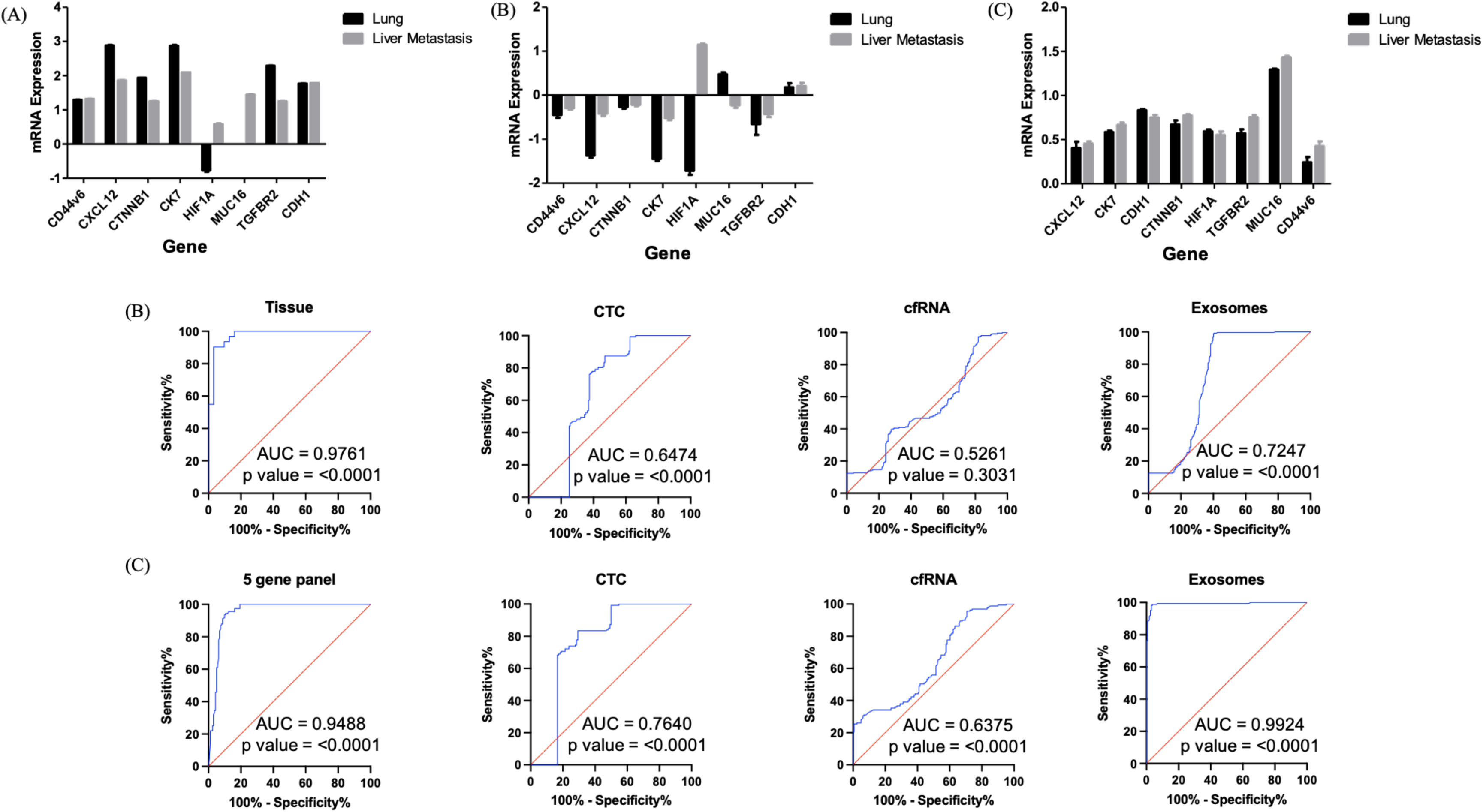

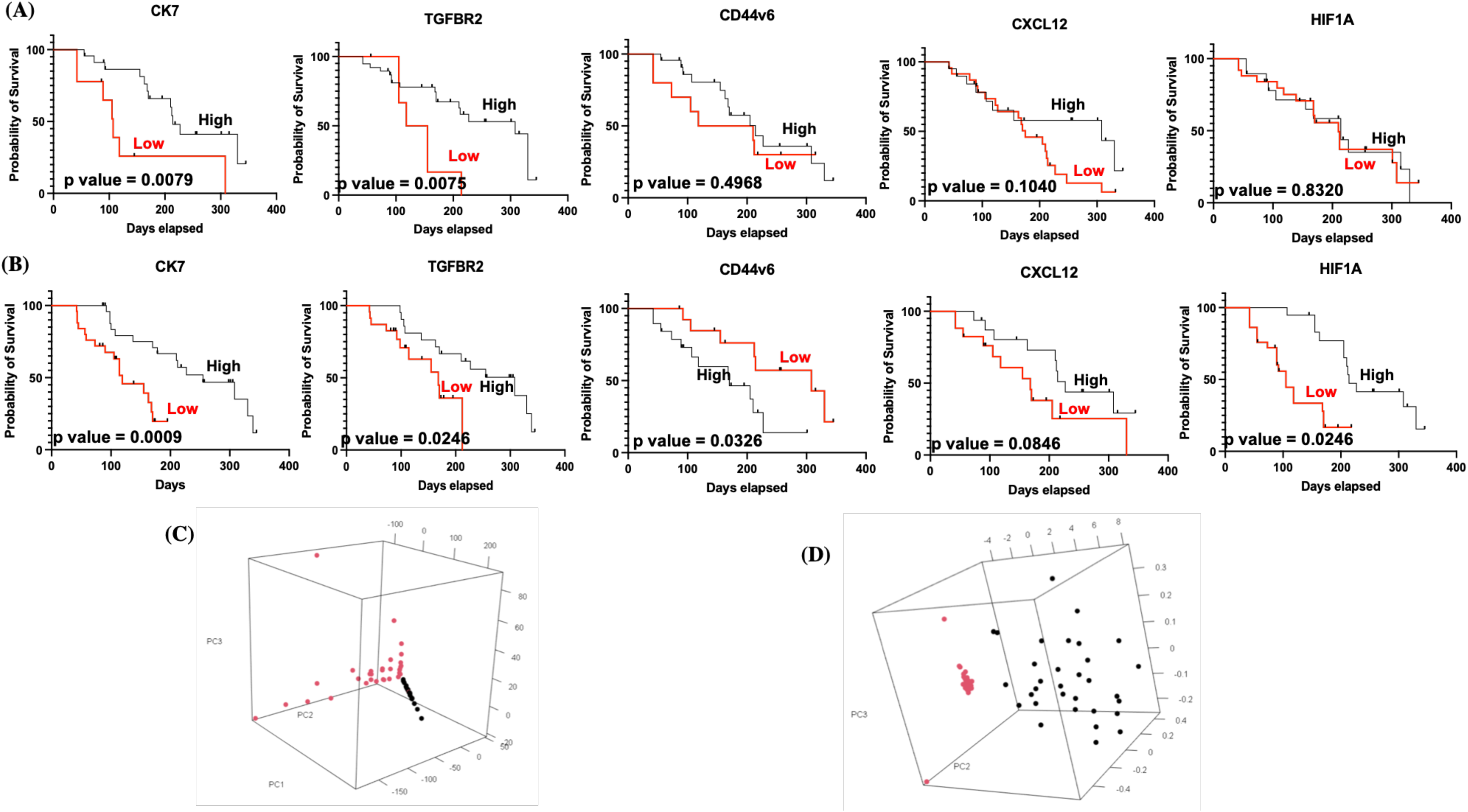

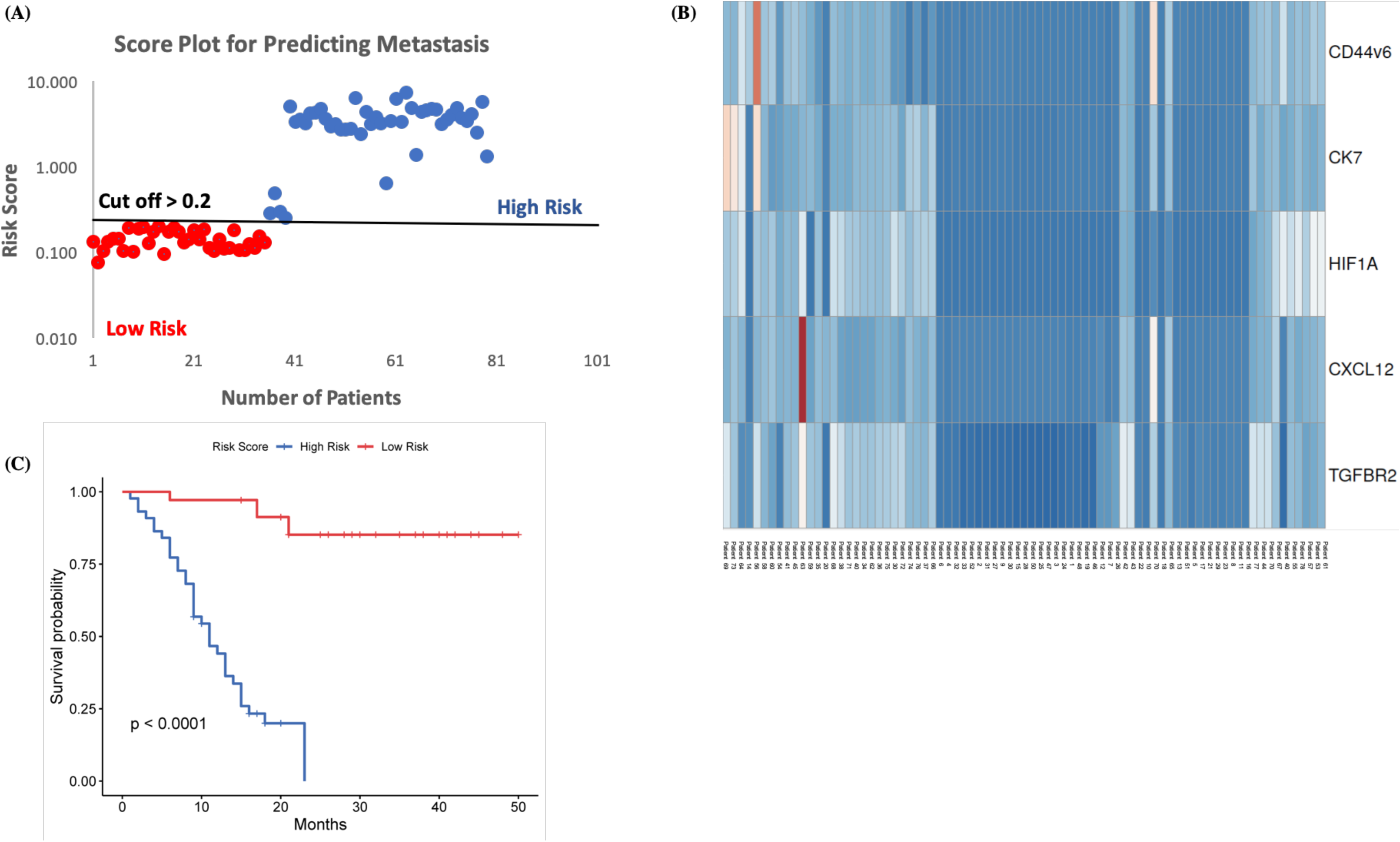

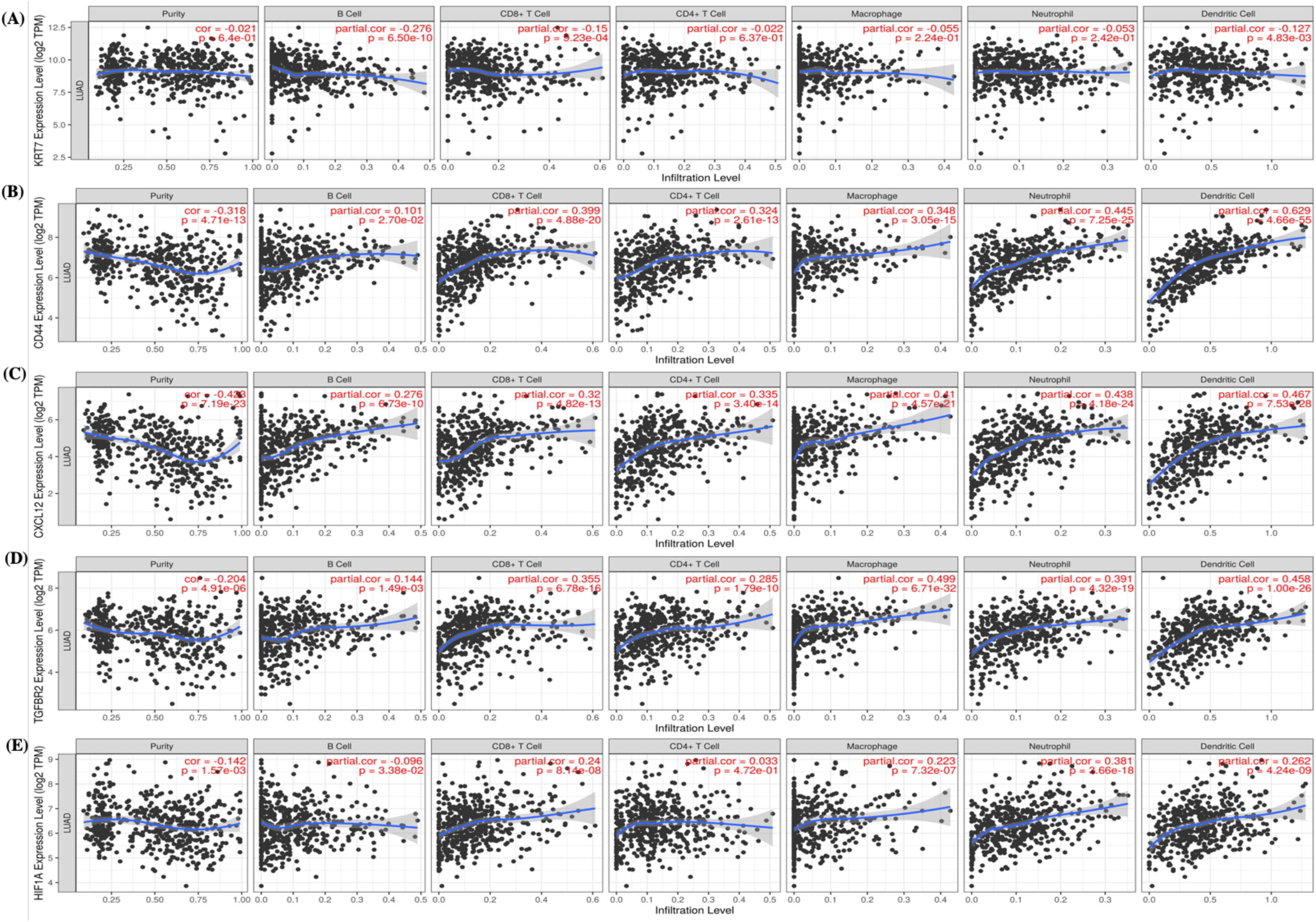

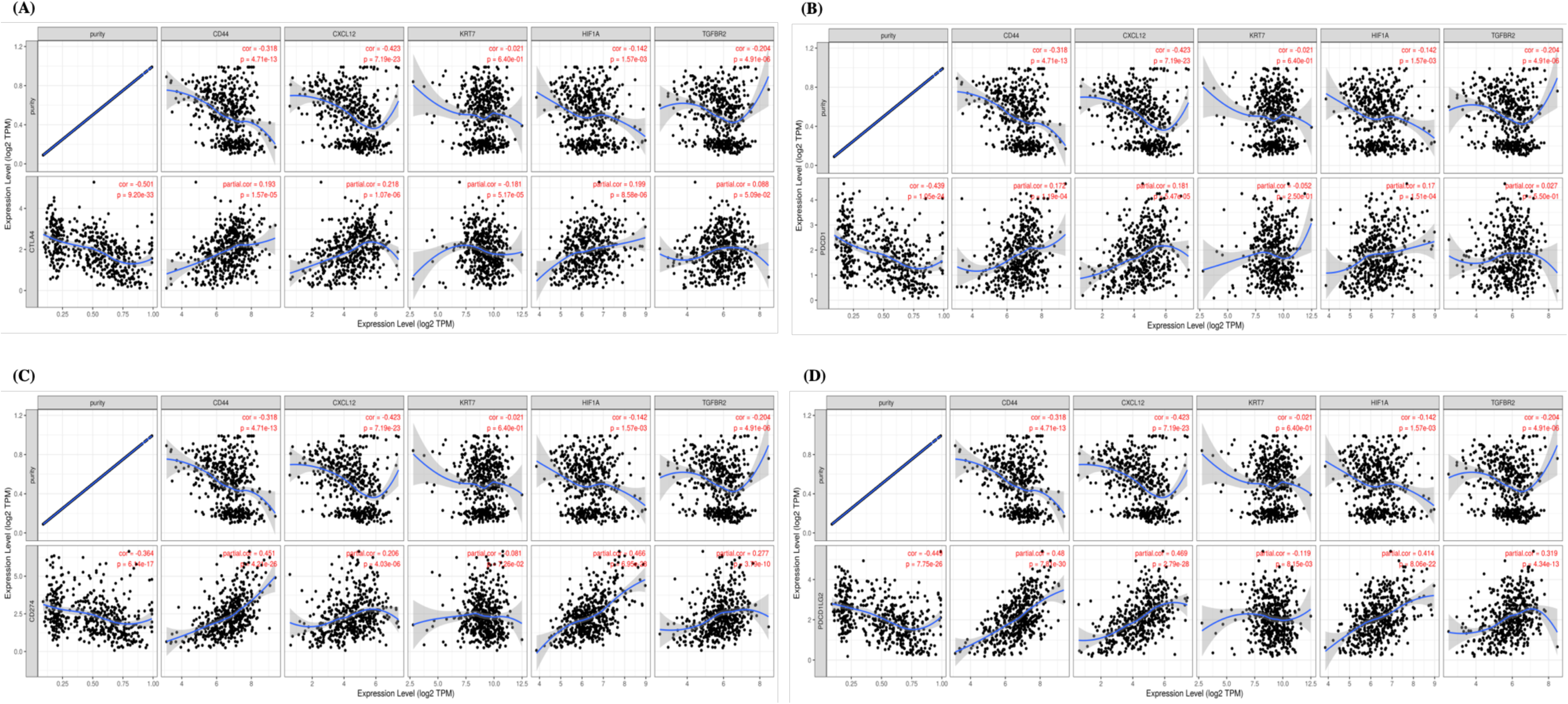

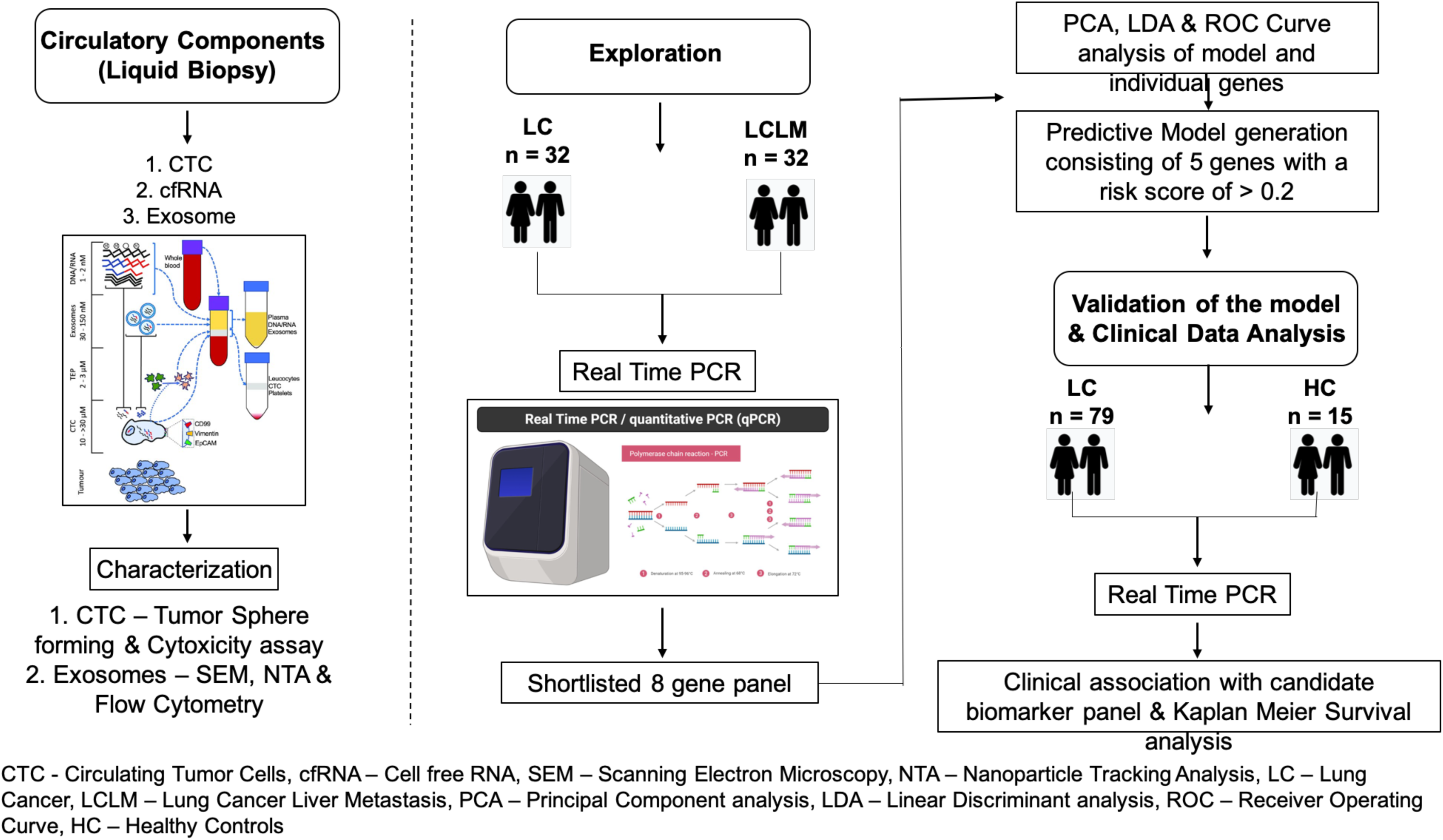

