## Supplementary figures and images for "Exosome derived multi-gene biomarker panel identifies the risk of liver metastasis in lung cancer patients"

### Supplementary Figure I

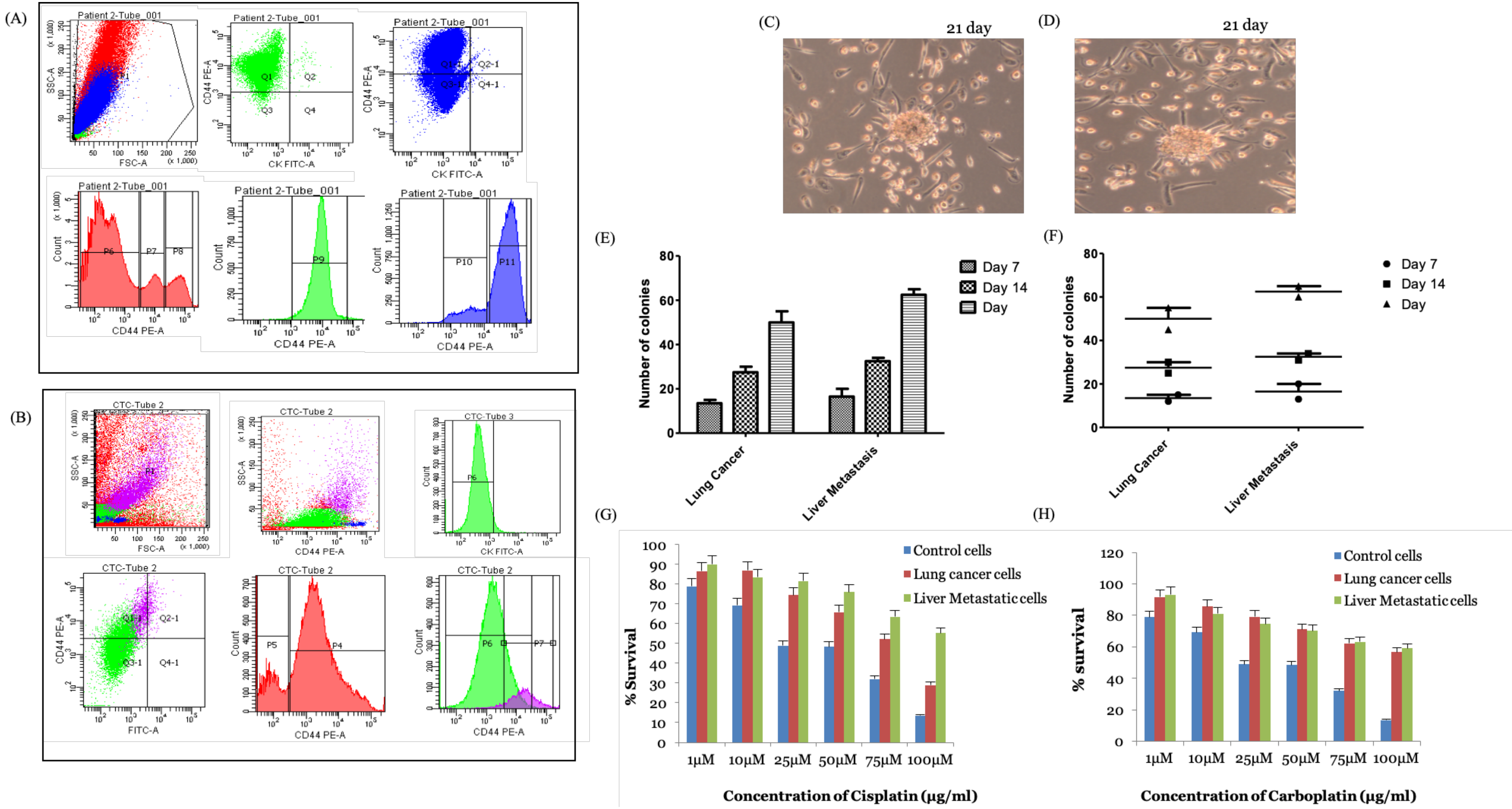

### Supplementary Figure III

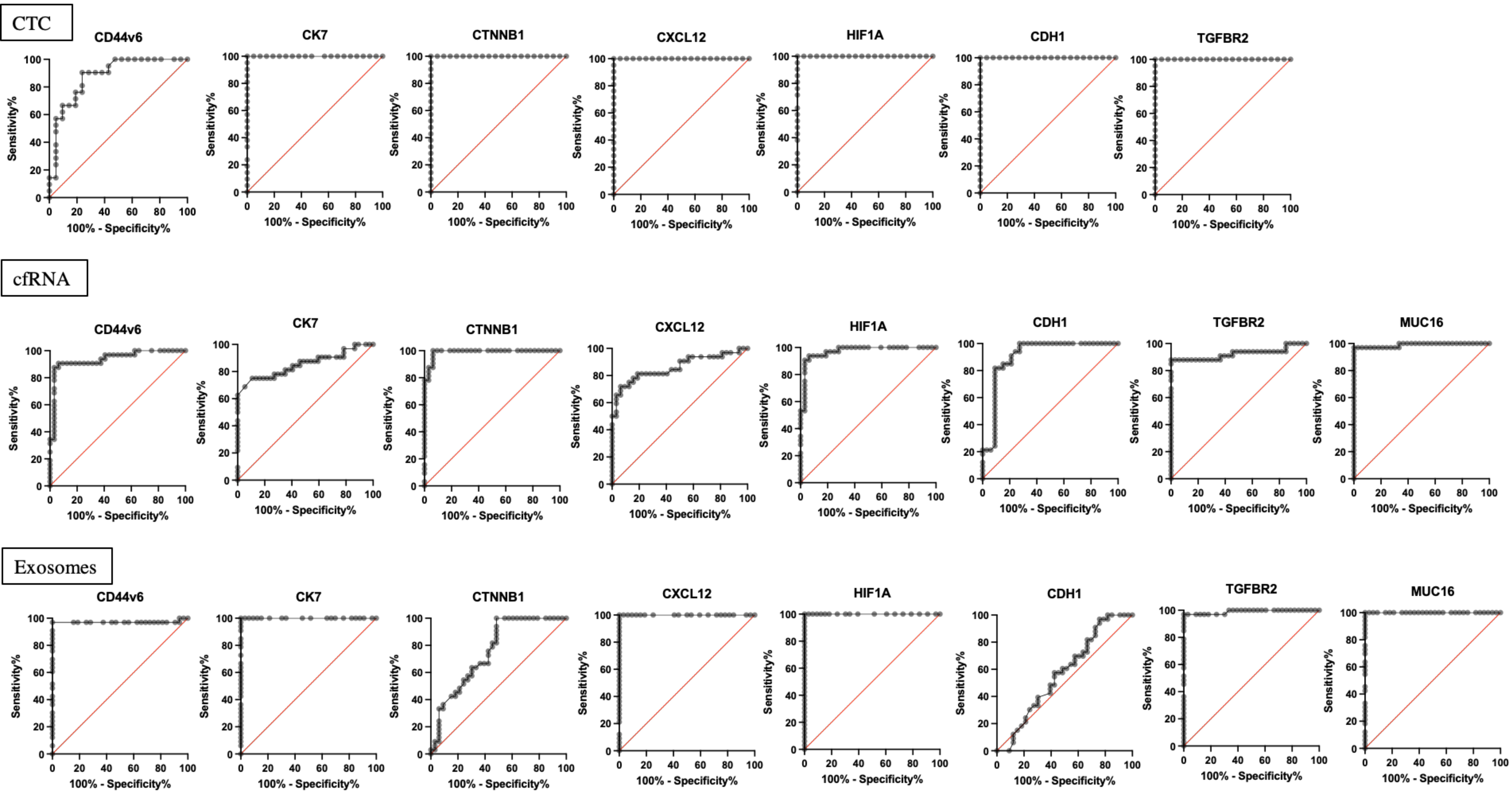
