## Supplementary Figure II for "Exosome derived multi-gene biomarker panel identifies the risk of liver metastasis in lung cancer patients"

(A)

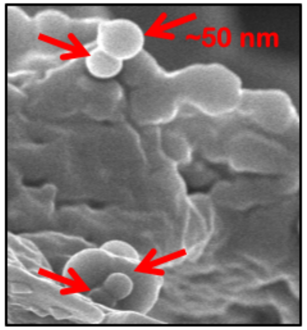


(B)

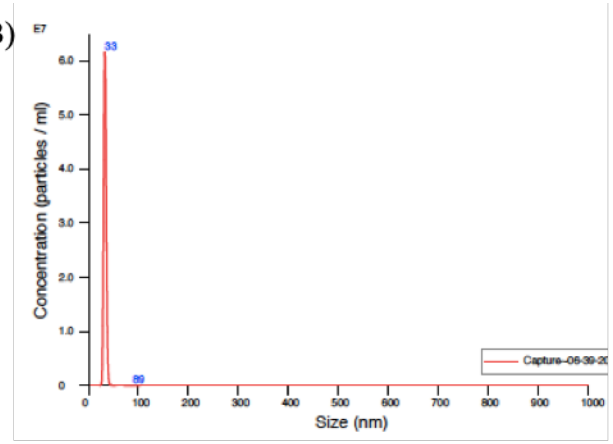


(C)

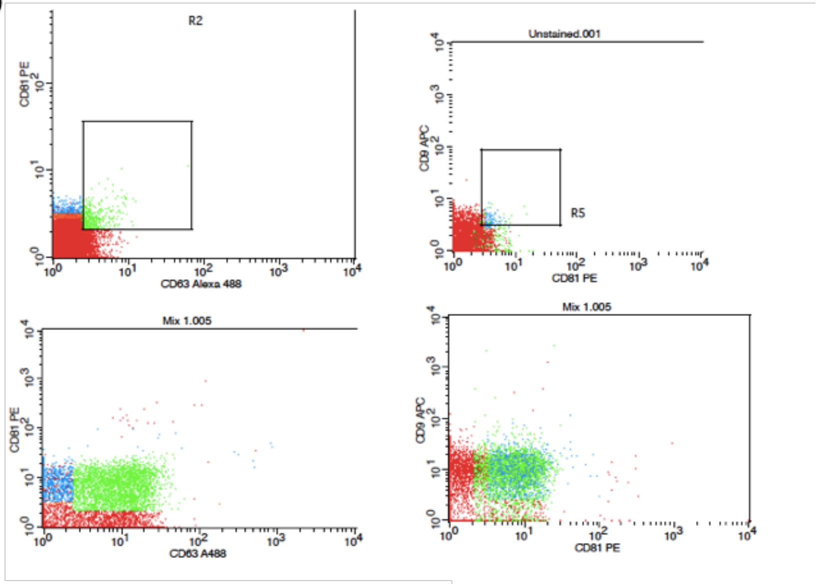


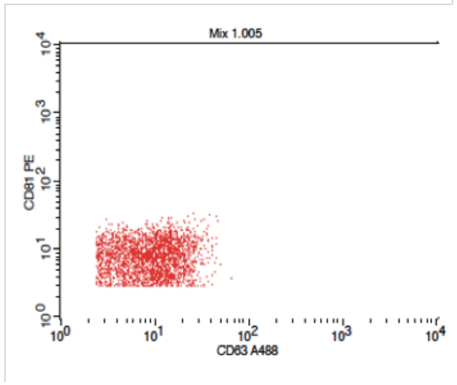


Gate Statistics

File: Mix 1.005
Sample ID: Mix 1
Tube: Untitled
Acquisition Date: 19-Sep-19
Gate: G1
Gated Events: 7876
Total Events: 10000

| Gate | Events | % Gated | % Total |
|---|---|---|---|
| G1 | 7876 | 100.00 | 78.76 |
| G2 | 3218 | 40.86 | 32.18 |
| G4 | 3623 | 46.00 | 36.23 |
| G5 | 3741 | 47.50 | 37.41 |
| G6 | 2482 | 31.51 | 24.82 |
| G7 | 3218 | 40.86 | 32.18 |
| G8 | 3741 | 47.50 | 37.41 |
